# Disentangling complex language contact and admixture in the broad Gansu-Qinghai region

**DOI:** 10.1101/2025.04.28.650932

**Authors:** Hongye Jin, Yuxin Tao, Chengkun Yang, Sizhe Yang, Wenjing Sun, Linguistic Silk Road Research Consortium, Dan Xu, Menghan Zhang

## Abstract

Language evolution in the Gansu-Qinghai (GQ) region provides a key perspective for understanding cultural development along the Silk Road. Previous genetic and archaeological studies revealed complex multiethnic interactions shaped by migration and sociocultural exchange. However, the lack of structured linguistic data and computational tools for studying language contact has limited rigorous analysis of sociocultural evolution in this area. Here, we presented a new hybrid dataset of phonological and morpho-syntactic features from languages sampled across the GQ region. We introduced a computational framework to assess language contact and admixture, allowing us to quantify interaction among GQ languages and trace their origins. Our results showed that GQ languages have distinct contact patterns in their phonological and morphosyntactic systems, with some languages exhibiting clear evidence of mixture. Using a new statistical method, Trait Sharing Test (*TS*-TEST), based on tree topology, we identified significant influences from Sinitic, Tibetan, Mongolic, and Turkic languages in shaping GQ linguistic diversity. These findings highlight the GQ region as a linguistic convergence zone on the Silk Road and provide a foundation for quantitative research on language contact and admixture. Our work strengthens the linguistic perspective on cultural evolution in the GQ region and supports future interdisciplinary studies integrating languages, genes, and material cultures.

## 1. Introduction

As a corridor linking Western and Eastern cultural spheres, the Silk Road has historically facilitated profound cultural, economic, and technological exchanges between China and Central/Western Asia ^1,2^. Positioned along the eastern section of this ancient trade network, the Gansu-Qinghai (GQ) region is notable for its extraordinary ethnic diversity ^3-5^, hosting communities such as the Han, Hui, Tibetan, and various Mongolic- and Turkic-speaking populations. It represents an enduring hub for multi-ethnic interaction and population admixture, spanning prehistoric and modern eras. Archaeological evidence indicates that at around 4,000 BP, populations cultivating wheat and barley from the west encountered millet-farming communities from the Loess Plateau in this region, leading to extensive technological exchanges ^6^. Genetic studies further demonstrate that the populations in the GQ region exhibit complex genetic compositions, with contributions from East Asia, the Tibetan Plateau, Siberia, and Central/Western Asia ^7-9^. Demographic studies also show that the GQ region remains a focal point for multiethnic migration and coexistence in contemporary times. ^10^ These all underscore its historical and modern significance as a cultural and genetic crossroads.

Language evolution is a bridge between demographic dynamics and sociocultural development. The interplay of population activities and socio-cultural interactions in the GQ region has given rise to high linguistic diversities, e.g., phylogenetic diversity and complex geographic patterns. Extensive efforts have been devoted to the linguistic fieldwork and documentation of languages in this region ^11-15^. Accordingly, several languages in the GQ region, such as Wutun ^16^ and Tangwang ^17^, have been categorized as “mixed” or “mixing” languages. Previous studies have also highlighted the GQ region as a linguistic area or *Sprachbund*, in which genetically unrelated languages share areal features due to subsequent contacts ^18-20^. In the GQ area, these features include word order, case-marking system, and strategies to express possession and presence ^19^. The susceptibility of these structural features to borrowing is consistent with established linguistic models^21^. When language contact is minimal, borrowing typically begins with non-basic vocabulary; As contact intensifies, structural features are increasingly affected. Structural borrowing starts with new phonemes with new phones in loanwords and shifts in the use of adpositions. It then extends to new phonemic contrasts and syllable structures in native vocabulary, as well as changes in word order and case-marking systems. Throughout this process, basic vocabulary remains little affected ^21^. Studies of language evolution primarily rely on basic vocabulary, as it shows less contact-induced changes ^22^ and provides divergence information. In contrast, structural features provide a more effective basis for investigating language contact and admixture, offering a lens for examining the interplay of language, population, and sociocultural dynamics in the GQ region ^22,23^.

Quantifying language contact and admixture presents a significant challenge. Over the past two decades, introducing computational methods inspired by evolutionary biology has provided new perspectives on the study of language evolution ^24-26^. Numerous biological approaches utilize genetic marker data to trace ancestral contributions within individual genomes^27^ or to analyze genetic admixture at the population level ^28,29^. These approaches can be adapted to linguistics by treating languages as analogous to individuals and linguistic features comparable to genetic markers ^30^. Several studies of language evolution have demonstrated the effectiveness of evolutionary biology methods in exploring linguistic structures and interactions. Examples include investigations into the languages of the ancient Sahul continent ^31^, the proposed Transeurasian language family ^23^, and Chinese dialects ^26^. Applying these methods offers a promising avenue for unraveling the GQ region’s complex language contact and admixture history.

Noting this, we aim to quantitatively disentangle language contact and admixture in the GQ region. Therefore, we compiled a dataset of 146 structural traits across 23 languages spoken geographically covering the GQ region (Figure 1). These languages can be classified into four linguistic phyla of northern China: Sinitic, Tibetan, Mongolic, and Turkic. We utilized a computational framework inspired by population genetics. This framework included linguistic relationships investigation, network analysis, population structure estimation, and admixture inference. Additionally, we developed a statistical test to quantify the influence of different sources on GQ languages. This test was based on comparisons of tree topology patterns, a method commonly employed to study introgression within evolutionary biology ^32,33^. By adopting this interdisciplinary approach, our study sheds light on the intricate multiethnic interactions that have shaped the GQ region.

**Figure 1.**
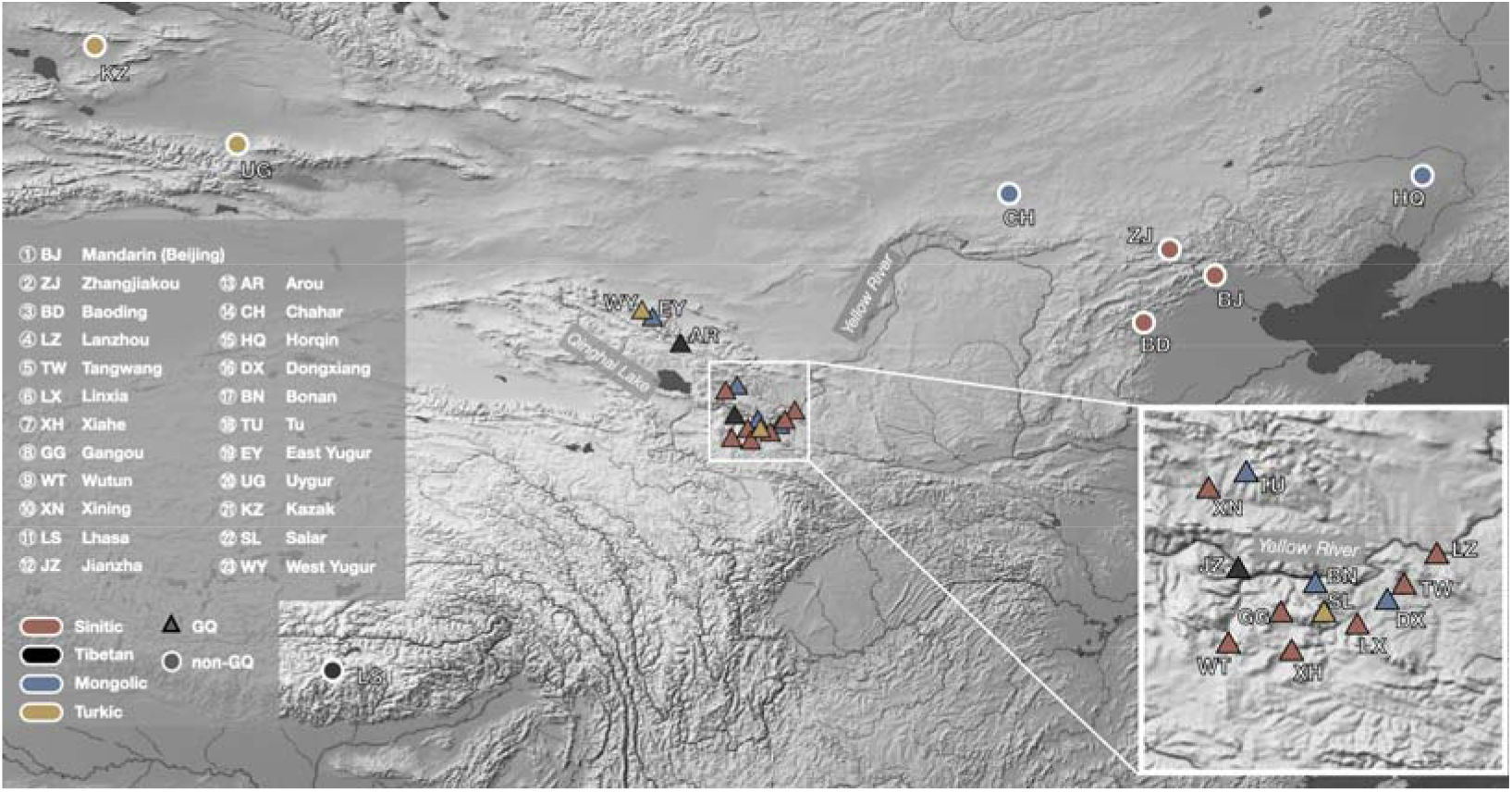
The 23 languages are included in this study. Colors indicate main phyla to which languages belong. Triangles with black borders represent languages in the GQ region, and circles with white borders represent non-GQ languages.

## 2. Results

### 2.1. Principal component analysis of linguistic structures in the GQ region

To investigate linguistic relationships among the 23 sampled languages, we conducted Principal Component Analysis using 146 structural traits of phonological and morpho-syntactic systems. The phonological PCA (Figure 2a) showed four primary clusters: (1) Sinitic languages and Lhasa Tibetan, (2) Amdo Tibetan dialects (Jianzha and Arou), (3) two Turkic languages (Uygur and Kazak), and (4) all Mongolic languages alongside with two Turkic in the GQ region (Salar and West Yugur). By comparison, the morpho-syntax PCA (Figure 2c) produced different clusters: (1) Sinitic varieties excluding Wutun, (2) Tibetan and Wutun, (3) Turkic languages with two GQ Mongolic languages (Bonan and East Yugur), and (4) other Mongolic languages. The key differences between the phonological and morpho-syntactic patterns were observed for Lhasa, Wutun, and four Altaic samples. Specifically, while Lhasa Tibetan and Wutun grouped with Sinitic in phonology, they aligned with Tibetan in morpho-syntax. Similarly, from the phonological perspective, Salar and West Yugur clustered with the Mongolic, while in morpho-syntax, Bonan and East Yugur clustered with the Turkic. Overall, the PCA results illuminated structural affinities within the linguistically diverse GQ region, offering insights into the complex interplay of phonological and morpho-syntactic characteristics.

**Figure 2.**
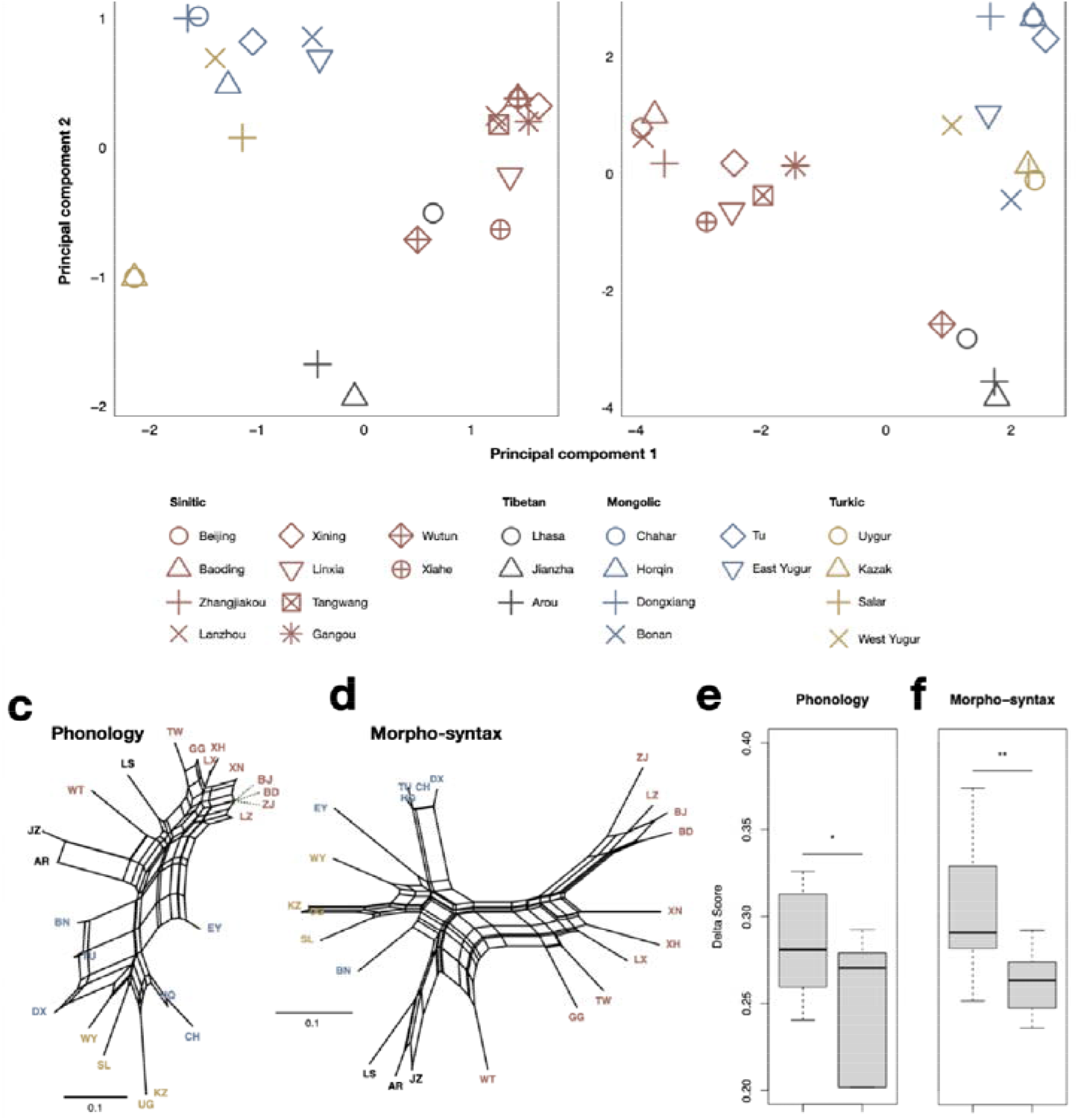
PC plots, Neighbor-nets, and delta scores of 23 languages. **a-b**: PC plots based on phonological (**a**) and morpho-syntactic (**b**) traits. **c**-**d**: Neighbor-nets based on pair-wise distances derived from phonological (**c**) and morpho-syntactic (**d**) traits. **e-f**: Phonological (**e**) and morpho-syntactic (**f**) delta scores for GQ and non-GQ languages. *p*-values were calculated using two-sample Wilcoxon test. *, *p*-value < 0.05; **, *p*-value < 0.01.

### 2.2. Neighbour-Net and delta score measurement of languages in the GQ region

To elucidate the interrelations among the 23 sampled languages, we calculated Hamming distances based on the presence or absence of phonological and morpho-syntactic traits, respectively. Pairwise distances were analyzed using the neighbor-net algorithm ^34^ (Figures 2c–d). Both networks exhibited extensive reticulations, highlighting widespread linguistic contact in both systems. In the phonological network (Figure 2c), Sinitic languages, East Yugur, Bonan, and Tu showed relatively short branch lengths and abundant box-like structures. On the other hand, the morpho-syntactic network (Figure 2d) showed the most intricate reticulations in GQ-Sinitic, East Yugur, West Yugur, and Bonan. These results suggested that these languages were involved in substantial horizontal influence.

To quantify the extent of language contact, we calculated delta scores ^35^ for each language. A two-sample Wilcoxon test comparing GQ and non-GQ languages showed that GQ languages had significantly higher delta scores in both phonological and morpho-syntactic systems (p < 0.05, Figure 2e-f), confirming more extensive contact within the GQ region. To assess whether contact dynamics in phonology and morpho-syntax were correlated, we computed Spearman and Kendall correlation coefficients for delta scores of GQ languages. Neither test showed statistical significance (Spearman: ρ = 0.257, *p*-value = 0.354; Kendall: τ = 0.143, *p*-value = 0.495), indicating no monotonic relationship between the two systems. These findings highlighted strong but distinct processes of language contact among GQ languages at the phonological and morpho-syntactic systems.

### 2.3. Fine-scale structure of languages from multiple ethnic groups

To investigate the fine-scale structure of the 23 languages, we used STRUCTURE to dissect their linguistic compositions based on phonological and morpho-syntactic traits, respectively. The *Evanno* method ^36^ favoured K = 3 as the optimal number of components for both analyses (Figure S1). In the phonological analysis (Figure 3a), the three components were Sinitic, Tibetan, and Altaic-related (blue, yellow, and green, respectively). In the GQ region, Sinitic languages were predominantly associated with the Sinitic-related component, except for Wutun, which displayed a significant Tibetan-related signal. Altaic languages in the GQ region showed varying similarities with East Yugur and Bonan, manifesting the highest proportions of Sinitic-related components. In the morpho-syntactic analysis (Figure 3b), the three major compositions were also Sinitic, Tibetan, and Altaic-related (red, gray, and orange, respectively). Within the GQ region, most Sinitic languages displayed notable Tibetan and Altaic-related components. The situation is particularly extreme for Wutun, which showed only a minimal proportion of the Sinitic-related component. Among Altaic languages in the GQ region, varying proportions of Sinitic and Tibetan-related components were observed. Bonan contained the highest proportion of Tibetan-related components, while East and West Yugur displayed considerable Sinitic-related components. Overall, these results shed light on classification and sharing patterns of structural features.

**Figure 3.**
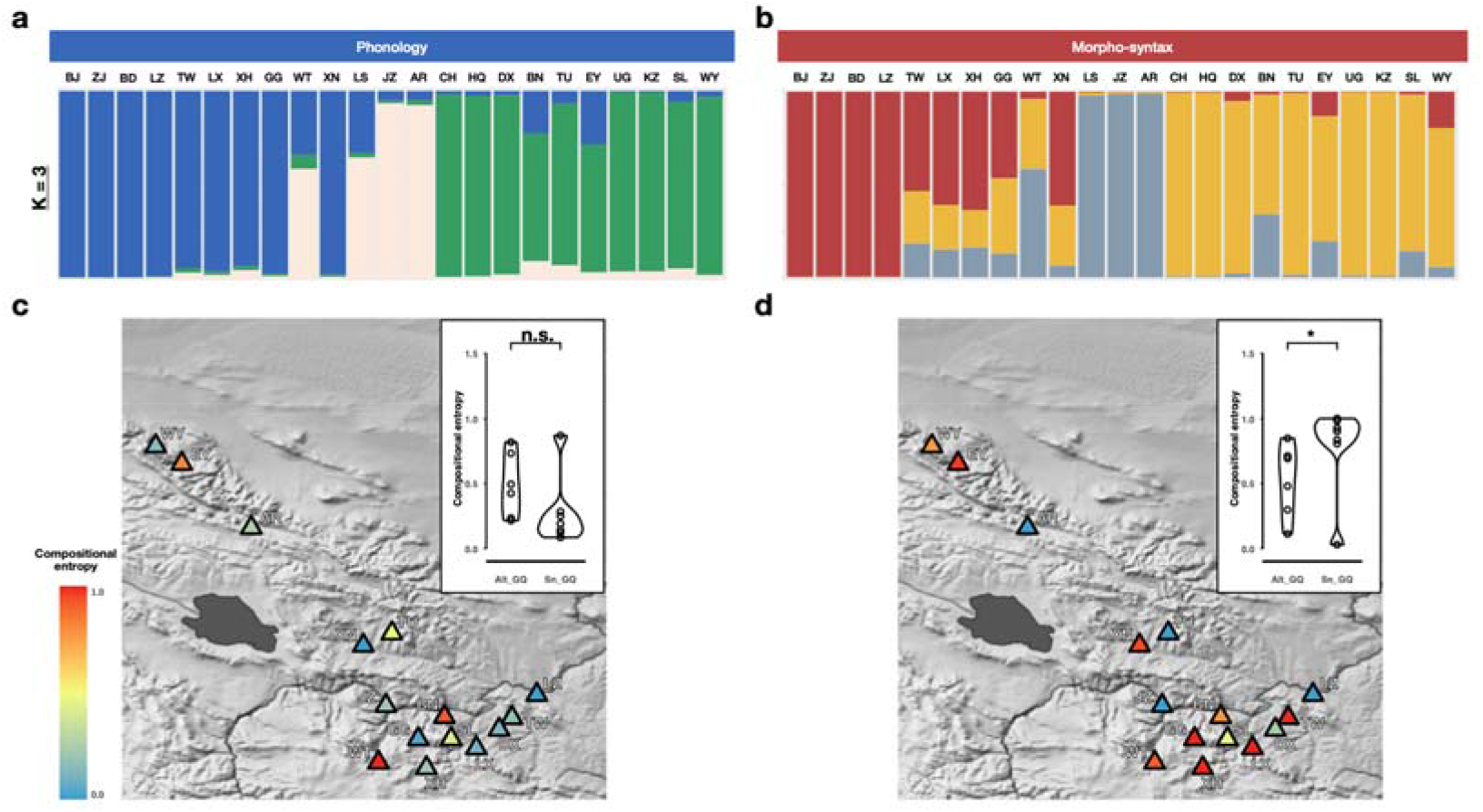
Fine-scale structure and compositional entropy of languages. **a-b**: Fine-scale structure of 23 languages inferred by STRUCTURE software using phonological (**a**) and morpho-syntactic (**b**) traits, respectively. **c-d**: Compositional entropy values derived from the K = 3 compositions of phonology (**c**) and morpho-syntax (**d**) for each GQ language. Upper-right windows show Wilcoxon tests comparing the GQ Altaic and GQ Sinitic entropy differences. n.s., not significant, *p*-value ≥ 0.05; *, *p*-value < 0.05.

To quantify the diversity of language components of GQ languages, we estimated compositional entropies for each language based on fine-scale structures at optimal K = 3 in both systems (Figure 3c-d). We observed that the compositional entropies of the two Amdo Tibetan dialects (Jianzha and Arou) were consistently low, while the patterns for Sinitic and Altaic languages in the GQ region were more complicated. We compared entropy values between GQ Sinitic and GQ Altaic languages and found that in morpho-syntax, GQ Sinitic have significant higher entropy values than GQ Altaic (two-sample Wilcoxon test, *p*-value < 0.05), while in phonology, GQ Altaic languages displayed higher mean and median entropy values than GQ Sinitic (mean: GQ Altaic = 0.46, GQ Sinitic = 0.31; median: GQ Altaic = 0.42, GQ Sinitic = 0.19), although the difference did not achieve statistical significance. These findings suggest that Amdo Tibetan in the GQ region has relatively simple structures in both systems, GQ Altaic languages exhibit more complex phonological structure of Altaic-, Sinitic-, and Tibetan-related components compared to GQ Sinitic languages, and GQ Sinitic languages show significantly more complex morpho-syntactic structure (See SI). In addition, we further examined the relationship between structural complexity in phonology and morpho-syntax across GQ languages based on their compositional entropy values. There was no statistically significant correlation (Spearman: ρ = -0.061, *p*-value = 0.832; Kendall: τ = -0.029, *p*-value = 0.923), suggesting independent evolutionary trajectories for phonological and morpho-syntactic complexity. Our results provide quantitative signals for language contact and potential admixture in the GQ region.

### 2.4. Assessing the linguistic contributions of neighboring languages to the GQ region

To examine whether the structural influences of GQ languages were from neighboring linguistic groups, we classified four language groups (GQ, Sinitic [Sn], Tibetan [Tb], and Altaic [Alt]) according to STRUCTURE results. Admixture *F3* statistics, computed from phonological and morpho-syntactic traits, showed significantly negative values for *F3(GQ; Sn, Alt)* (Figure 4a; phonology: p = 0; morpho-syntax: p = 0.042; bootstrap permutation test; see Table S1-2 for full results). This indicated a mixture of Sinitic and Altaic influences on the GQ language group. To corroborate this finding, Bayesian network modeling further identified directed edges from Sn, Alt, and Tb language groups to the GQ group (Figure 4b, S9; see Table S4-5 for full results), highlighting their contributions. These findings demonstrate that combined influences from Sinitic, Altaic, and Tibetic languages have shaped the structural features of languages in the GQ region.

**Figure 4.**
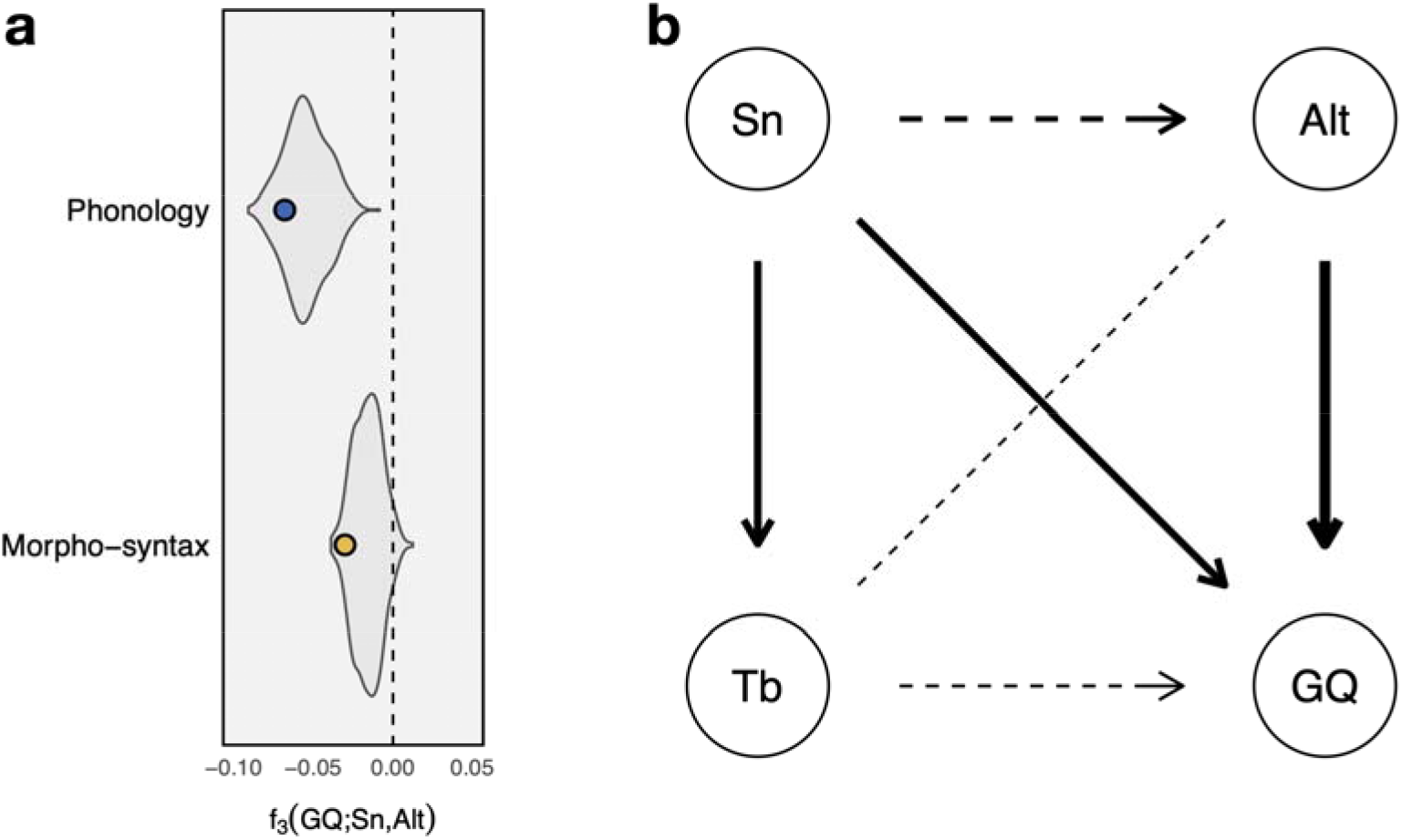
Admixture statistics and Bayesian network. **a**: Dots present admixture statistics measured using morpho-morpho-syntactic and phonological traits. The violin plots show distributions derived from bootstrap permutations. **b**: A summary plot of Bayesian network results, adapted from the network constructed using overall data, integrating results from phonological and morpho-syntactic analyses. Solid arrows indicate edges that appeared in the analysis using overall data; dashed arrows indicate the edge did not appear in the analysis using overall data but appeared in the phonological or morpho-syntactic analysis. The dashed line without arrows indicates conflicting results across the three analyses (See SI). The thickness of the arrows is related to the strength of edge importance.

### 2.5. Inferring admixture sources on individual GQ languages

To investigate the influence of four major language groups in northern China—Sinitic, Tibetan, Mongolic, and Turkic—on languages in the GQ region, we developed a statistical method, the Trait Sharing Test (*TS*-Test). Based on tree topology comparisons, this method assesses whether the trait sharing between a target language (a language in the GQ region) and a source language group (one of the four) is statistically significant. A significant result indicates a substantial contribution from the source group. Phonological and morpho-syntactic traits were analyzed separately.

Based on the *TS*-Test, the phonological analysis (Figure 5a) revealed that most GQ languages primarily reflected contributions from their respective phyla. For example, Xining, a Sinitic language, exhibited substantial contributions from the Sinitic group. West Yugur showed significant input from both Turkic and Mongolic sources. We found strong Sinitic influence in East Yugur. Wutun had pronounced influences from both Tibetan and Sinitic. In contrast, Salar displayed no statistically significant evidence of phonological trait sharing with any of the four groups. This may be due to minor or diffuse contributions from multiple sources, as reflected in its fine-scale structure (Figure 3a), which resulted in a random-like pattern. Therefore, the small number of phonological traits in our dataset (n = 26) may have limited the resolution necessary to detect these influences. On the other hand, the morpho-syntactic results (Figure 5b) revealed more complex patterns of language admixture. West Yugur, East Yugur, Tu, Salar, Wutun, and Gangou exhibited substantial contributions from two source groups. Bonan demonstrated influence by the greatest number of sources, receiving significant contributions from three groups: Tibetan, Mongolic, and Turkic. These findings provided a detailed perspective on how linguistic interactions with neighboring groups have shaped individual GQ languages.

**Figure 5.**
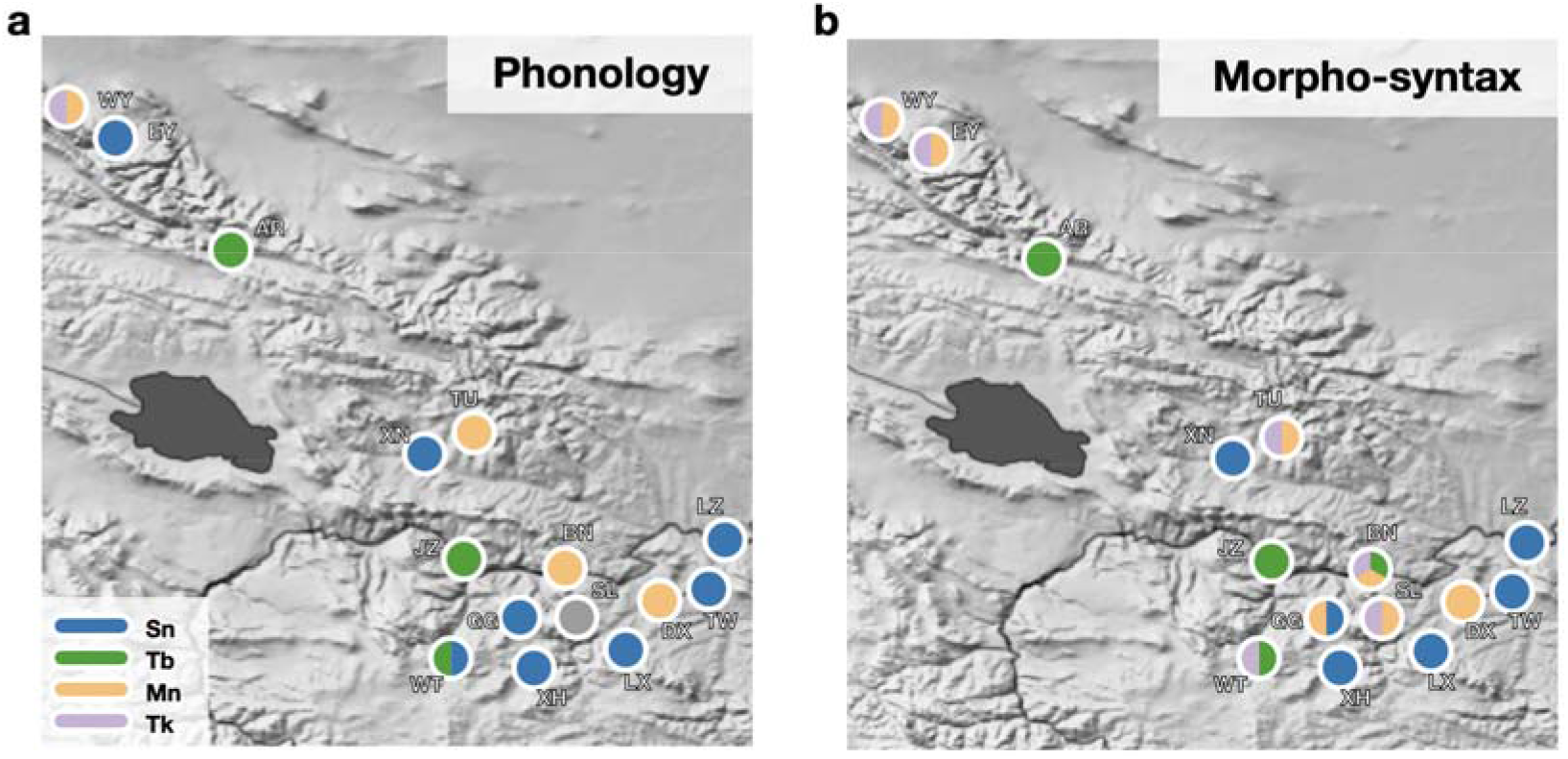
Significant trait sharing between languages in the GQ region and the four sources. The segments in the pie charts represent substantial contributions from sources corresponding to respective colors. A gray circle for Salar in the phonological plot indicates that it showed no significant influence from any of the sources.

## 3. Discussion

The Silk Road was one of the most transformative exchange networks in human history, facilitating the convergence of diverse peoples, goods, and cultures. Positioned along the eastern stretch of this historic corridor, the Gansu-Qinghai (GQ) region has been a critical nexus of multi-dimensional interactions that have profoundly shaped the cultural and linguistic landscape of Northwest China. Understanding the patterns of multi-ethnic contact in this region is central to exploring its rich sociolinguistic complexity. In this study, we compiled a dataset of 146 structural traits of phonological and morphosyntactic systems from 23 languages, which offered robust material for examining linguistic admixture in the GQ region. According to the dataset, we proposed a new computational framework to quantitatively disentangle the language contact and admixture. Our findings revealed that linguistic interactions in the GQ region exhibited high intensity, with distinct patterns governing phonological and morphosyntactic systems. By applying population genetic approaches, we inferred the fine-scale linguistic structure of these languages and demonstrated that the observed structural patterns in the GQ region reflect the combined influence of Sinitic, Altaic, and Tibetan language groups. Additionally, we developed a statistical tool to test trait sharing and showed how Sinitic, Tibetan, Mongolic, and Turkic sources have shaped the structural diversity of GQ languages.

The GQ region is characterized by its striking linguistic diversity, with different languages from distinct phyla coexisting in proximity. Our PCA of 23 language samples identified four major clusters in both phonological and morpho-syntactic systems, which broadly corresponded to the Sinitic, Amdo Tibetan, Turkic, and Mongolic language groups. Language contacts among these groups led to the complex process of admixture, as exemplified by Wutun and four Altaic languages spoken locally (Bonan, East Yugur, Salar, and West Yugur). Wutun is linguistically recognized as a mixed language ^16^. Its grammatical structure has been profoundly shaped by Amdo Tibetan, meanwhile, most of its lexicons derive from Mandarin Chinese, which results in a phonological system compatible with other Chinese dialects ^37^. Consistent with these descriptions, our PCA showed Wutun clustering with Sinitic languages in phonology, while with Amdo Tibetan in morpho-syntax. Among Altaic languages in the GQ region, previous linguistic studies proposed that East and West Yugur form a “mini-*Sprachbund*,” with Bonan and Salar constituting another ^20^. In this region, several phonological features, such as consonant strength and specific phonemic merging ^20^, were shared by Mongolic and Turkic languages. Additionally, morpho-syntactic traits in GQ Mongolic, such as plural markers and conditional converbs, appear to reflect Turkic impact ^14^. Our findings in this study support these observations as Turkic languages (Salar, West Yugur) grouped with Mongolic languages in the phonological PC space, while Bonan and East Yugur (Mongolic) clustered with Turkic languages in morpho-syntax. These patterns highlighted language contact and admixture among Mongolic and Turkic languages in the GQ region. Moreover, GQ Sinitic languages exhibited distinctive areal features compared to other Chinese varieties, such as OV word order and case marking^19^, resulting from contacts with non-Sinitic groups. Our morpho-syntactic PCA exhibited a similar pattern in which five GQ Sinitic languages clustered close to each other. These results aligned with previous studies and strengthened the evidence for GQ as a linguistic area.

Different language systems followed different dynamics during language contact and admixture in the GQ region. Previous studies report that Mongolic and Turkic languages in this region experienced varying degrees of Tibetanization or Sinicization in their phonological systems ^38^. At the same time, GQ Sinitic varieties extensively borrow morpho-syntactic features from Altaic and Tibetan languages ^18-20^. Our computational analyses provided new evidence supporting these patterns. We found that the intensities of phonological and morpho-syntactic contacts and admixture in GQ languages should be largely independent. Specifically, GQ Altaic languages showed more complex phonological strcuture than GQ Sinitic, whereas GQ Sinitic languages exhibited significantly more complex morpho-syntactic structure.

Notably, language contact and admixture in the GQ region can be partially explained by the genetic history of its populations. None of the groups speaking GQ languages are genetically homogeneous ^19^. For example, speakers of Bonan and Salar prevalently carry Y-haplogroups associated with Sino-Tibetan populations ^9,39^, and we showed that both languages contain notable Sinitic and Tibetan-related components. However, some cases in the GQ region illustrate a mismatch between linguistic affiliation and population activities. The ancestors of Bonan, Dongxiang, and Salar speakers likely originated from Turkic-speaking populations in Central or Western Asia ^9,40,41^; however, Bonan and Dongxiang peoples shifted to Mongolic languages, while Salar people retained their Turkic language. Other examples include the Wutun speakers. Linguists have previously hypothesized that ancestors of Wutun speakers may have migrated from Nanjing or Sichuan ^42^. Currently, Wutun bears strong Amdo Tibetan influence, and its speakers self-identify as Tibetan. Overall, detailed genetic studies of speakers of languages in the GQ region remain limited. Current genetic research has primarily focused on large-scale east–west admixture patterns, and the genetic background of each population in this region still requires further investigations. More comprehensive interdisciplinary research is needed to clarify the complex interplay of languages, genes, and cultures in the GQ region.

In the GQ region, certain linguistic structural features have played key roles in the dynamics of language contact and admixture and have been shaped by intensified selective pressures. Among ten structural traits identified in our study (Table S7), case-marking systems were particularly noteworthy, including *AccDat* (identical accusative and dative case markers), *ErgInst* (identical ergative and instrumental markers), and *NomAcc* (nominative-accusative alignment). The adoption of case marking in Sinitic languages is a defining characteristic of the GQ linguistic area ^18-20^. A previous study, based on an early 20th-century Gansu dialect corpus, suggested that case markers in GQ Sinitic originated from prosodic fillers used as topic markers, arising through sustained contact between Sinitic and non-Sinitic speakers ^43^. Our findings echoed this view, highlighting the pivotal role of case-marking features in language contacts in the GQ region. In addition, several traits were instrumental in distinguishing the three linguistic components identified in the region’s fine-scale language structure ^23^ (See Materials and methods and Figure S2). For example, vowel harmony in the past (*HarmVPast*) distinguishes the Altaic-related component in phonology, while prepositions and postpositions (*PrepPost*) mark the Sinitic-related component in morpho-syntax. These findings provided data-driven analyses of crucial features during language contact and admixture in the GQ region.

In addition, we employed tree topology comparisons to disentangle the influences of distinct source languages on target languages and introduced a statistical approach, *TS*-Test. This approach assesses the proportion of shared traits between targets and sources while controlling similarities arising by chance through statistical evaluation. As the number of traits increases, we expect the accuracy of this method to improve accordingly. Applying this approach, we delineated the respective contributions of Sinitic, Tibetan, Mongolic, and Turkic languages to those spoken in the GQ region. We anticipate that *TS*-Test will facilitate broader culture-related quantitative investigations, offering a useful tool for exploring linguistic and cultural exchanges across diverse contexts.

## 4. Conclusions

In this study, by analyzing structural features across 23 languages, we confirm extensive language contact and admixture in the GQ region, supporting previous proposals of GQ region as a linguistic *Sprachbund* ^18-20^. Our results reveal distinct dynamics of contact and admixture in phonological and morphosyntactic systems. The overall structural pattern of GQ languages can largely be attributed to the combined influences of Sinitic, Mongolic, Turkic, and Tibetan languages (Figure 6). However, the extent of each influence varies across individual GQ languages. Key structural features, such as case-marking strategies, highlight the role of multi-ethnic contact in shaping the linguistic landscape of this region. Future studies should expand the number of language samples and linguistic features to better understand the GQ region, Northwestern China, and the Silk Road. Integrating linguistic, genetic, and archaeological evidence will illuminate the intricate history of multi-ethnic exchanges along the Silk Road.

**Figure 6.**
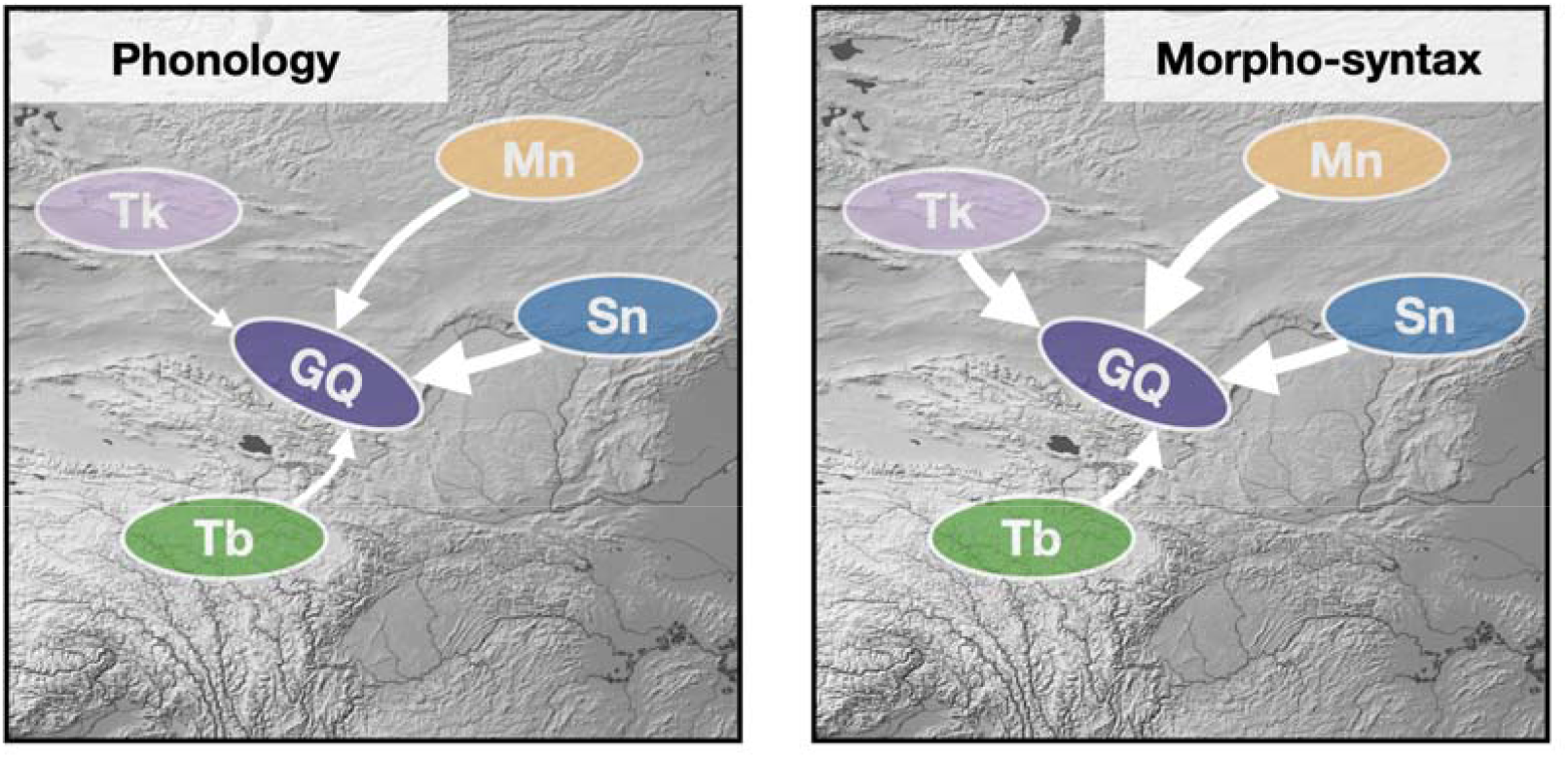
Summary of linguistic contact and admixture in the GQ region as influenced by surrounding linguistic groups.

## 5. Materials and methods

### 5.1. Data Collection

The linguistic data were collected by linguists working in different languages. Among the 23 language datasets, 18 were obtained from recent fieldwork, and 5 were sourced from existing documents. A list of these authors is annexed at the end of this paper. The dataset contained 26 phonological traits and 120 morpho-syntactic traits across 23 languages. We designated this dataset as GQ_admix. In this dataset, there are 15 languages sampled from the GQ region. These comprised seven Sinitic varieties (Xining, Linxia, Tangwang, Gangou, Wutun, Xiahe, and Lanzhou), two Amdo Tibetan (Jianzha and Arou), four Mongolic (Dongxiang, Bonan, Tu, and East Yugur), and two Turkic (Salar and West Yugur). Besides, eight languages outside the GQ area are included—Standard Mandarin (Beijing), Baoding, Zhangjiakou, Lhasa, Chahar, Horqin, Uygur, and Kazak.

### 5.2. PCA, Neighbor-net, and delta score

To give profiles of the 23 languages, we conducted principal component analysis (PCA) based on their structural traits. The PCA was performed using the prcomp() function from R *base* package, and the proportion of variance explained by each principal component was calculated using the fviz_eig() function from the *factoextra* package. Next, we implemented the Neighbor-net algorithm ^34^ to further examine language contact. Neighbor-Net is a phylogenetic network model based on an agglomerative method. A higher number of box-like structures in a derived network indicates greater horizontal exchange. To quantify the contact strength of languages, we calculated the delta scores ^35^ for each language. The delta score is a statistical measure for detecting non-treelike signals in data; a higher delta score indicates that a language has obscured more treelike signals and suggests stronger contact. These methods have been widely used to assess tree-like patterns in linguistic data ^22,26,44^. The Neighbor-net was constructed using R package *phangorn*. Delta scores were computed using SplitsTree v6.3.35 ^45^. We specified Hamming distance as the distance measure.

### 5.3 Fine-scale Structure of Languages

To investigate the nuanced structure of languages, we conducted population stratification analysis using the software STRUCTURE v2.3.4 ^27^. Following the similar computational process in the previous studies ^23,26,31^, we performed separate clustering analyses on phonological and morpho-syntactic data, respectively. Considering the nature of linguistic data, we set the ploidy level of language individuals to 1. We employed the “admixture” model and allowed the model to estimate the Dirichlet parameter α for admixture proportions. We allowed independent allele frequencies across populations and fixed the Allele Frequencies Parameter λ At 1.0. We made the STRUCTURE software calculate from the hypothetical population number K = 2 to K = 10, and for each K value, we performed 30 repetitions. We set the Markov chain to run for 10,000 generations after 10,000 generations of burn-in.

We analyzed STRUCTURE outputs using structureHarvester ^46^, which implemented the *Evanno* method ^36^ to calculate ΔK statistics based on the rate of change in log probability between successive K-values. The hypothetical number corresponding to the highest ΔK values represents the optimal number of components. Replicates for each K-value were summarized using the Greedy algorithm in CLUMPP ^47^. Results were visualized through stacked bar plots.

To quantify the diversity of language components, we calculated the compositional entropy using Shannon entropy estimation for each language based on the fine-scale structure inferred at the optimal K number. Higher entropy values indicate a higher linguistic diversity of languages. We calculated such entropies using the R package *entropy*. We performed two-sample Wilcoxon tests to assess differences in linguistic diversity of languages between GQ Sinitic and GQ Altaic.

To identify instrumental traits in delineating components, we examined inferred allele frequencies for each component provided in STRUCTURE outputs. We selected the replicate with the highest Estimated Prob of Data at optimal K in phonological and morpho-syntactic analyses, respectively. For each hypothetical component, we manually extracted the frequency of each trait being present (value = 1). Traits that were present (frequency > 2/3) in at least one component and were absent (frequency < 2/3) in one other component were selected as critical traits.

### 5.4. F3 Statistics and Bayesian Networks

To investigate whether the admixture of surrounding language groups can explain the structural features in the GQ region, we employed two methods: the admixture *F3* statistic and the Bayesian network.

The admixture *F3* statistic, introduced by Patterson ^28^, is a widely adopted measure in population genetics for assessing whether a population represents an admixture between two other populations. In the genetic analysis, *F3(C; A, B)* is defined as the average product of allele frequency differences at each genetic marker between a target population C and two source populations A and B. A negative *F3* value indicates that the target population C is a mix of populations A and B.

Analogously, in our analysis, we treated linguistic structural traits as genetic markers and language groups as populations. We defined four language groups (GQ, Sinitic [Sn], Tibetan [Tb], and Altaic [Alt]) based on STRUCTURE results. These groups included languages that maximally represent each of the three inferred components and potentially mixed GQ languages. Notably, although Dongxiang was constituted by a relatively pure Altaic-related component in both systems, we did not include it in the Alt language group but in the GQ language group. This was based on the fact that Dongxiang is located deep in the GQ linguistic area geographically, and it was reported to have some highly distinctive characteristics, making it not readily intelligible to any other Mongolic language speakers ^14^. We then computed *F3(GQ; Sn, Alt)*, *F3(GQ; Sn, Tb)*, and *F3(GQ; Tb, Alt)* using phonological, and morpho-syntactic traits in our dataset. The computational procedure involved two main steps. First, we calculated trait frequencies within each language group. For each trait, we computed the product of frequency differences between GQ and each pair of source groups. The *F3* statistic was then derived as the means of these products across all traits. Second, to assess whether the obtained *F3* values were significantly negative, we utilized a permutation test framework. We performed a bootstrap resampling process with 500 iterations. A value was considered significantly negative if more than 95% of bootstrap replicates yielded negative values. The distribution of bootstrap results was visualized using a violin plot. All *F3* calculations were implemented through a custom R script.

Bayesian networks are directed acyclic graph models that represent a set of variables and their conditional dependencies ^48^. In a Bayesian network, nodes are sets of variables, and edges (arrows) between nodes indicate direct probabilistic dependencies. The state of a downstream node depends on the state of an upstream node. In the current study, we interpreted this as indicating a directed contribution from one language group to another. We constructed Bayesian networks using structural trait frequencies from the four language groups (i.e., GQ, Sn, Tb, Alt) and employed a penalized node-average likelihood approach ^49^ for network scoring. The model determined the presence and directionality of arrows between nodes, yielding an optimal network structure, and assessed the importance of each connection. To ensure comparability across networks, we reported each edge’s importance as its proportional contribution to the overall model likelihood. We interpreted this value as the relative influence of an upstream node on a downstream node. The Bayesian networks were constructed using the hc() function from the R package *bnlearn*, and the importance of each connection was scored using the package’s score() function.

### 5.5 Language trait sharing testing

To quantify the trait sharing between a target language and multiple source languages, we derived a statistical method based on the comparison of tree topology patterns. We named this tool Trait Sharing Test (*TS*-Test). This method used triadic combinations as the basic analytical unit (See SI). Given a target language *t* and a set of source languages *S* (with at least two sources), the system iterated through all possible triadic combinations formed by the target and different source pairs {*t, s_i_, s_j_J*. In the first step, within each triadic combination, the analysis first filtered traits by removing those with missing values and eliminating invariant traits (where t, s_i_, and s_j_ share identical values), resulting in a subset of valid traits. Subsequently, each trait can be used to construct a tree involving the target and two sources, whose topology indicates whether the target aligns with one of the sources or is distinct from both. It then calculated the matching rate between the target and each source language, i.e., the proportion of trees with the target and a source clustered together. In the second step, at the global level, each source language’s local matching rates across all its participating triadic combinations were averaged, and the results were normalized to produce a global sharing score □_∈_ [0,1].

We implemented a permutation test framework to assess the statistical significance of the trait sharing between target and source languages. We produced 5000 permutation samples of each target language’s binary traits vector while maintaining the marginal traits counts. For instance, if the actual data contained 50 traits with 20 taking value 1 and the remaining 30 traits taking value 0, the permuted samples would preserve this 20:30 distribution. This process generated a null distribution of similarity scores, from which we constructed the 95% Highest Density Area (HDR). If the observed sharing score □ exceeded the upper threshold of the corresponding 95% HDR, we would reject the null hypothesis that the target and source language trait distributions were independent, concluding that their trait sharing was statistically significant.

In *TS*-Test, both the target and source can represent individual languages or language groups. If a target or source contains multiple languages, the program uses vector replication during triadic combination calculations to ensure all possible inter-language combinations are considered. Specifically, if a triadic combination contains two source groups with *m* and *n* languages, respectively, and a target group with k languages, the trait vectors of languages in these source groups will be replicated *n × k* and *m × k* times respectively. In contrast, the target language vectors will be replicated *m × n* times. This expanded vector set will contain *m × n × k* combination configurations.

We represented the observed sharing scores using stacked bar plots and showed the null distributions of different sources using stacked kernel density plots. Sources showing statistically significant trait sharing were marked with asterisks. The R scripts required to execute the *TS*-Test are provided in the supplementary files.

### 5.6. Outlier Traits Detection

To better understand the traits that are crucial in shaping the scenario of languages, we conducted a PCA-based outlier detection to identify outlier traits showing distinct distributions compared to most traits. We conducted outlier detection separately using phonological, morpho-syntactic, and overall traits in our dataset. We excluded traits with variant frequencies below 5%. After PCA, we selected the top three principal components for the following use, as they collectively explained over 50% of the variance in all three analyses. This method performed regression between traits and principal components, generating z-scores that indicate the degree of deviation from the overall mean. Based on these z-scores, it calculated distance statistics that should follow a chi-square distribution under the null hypothesis of no outlier traits, thereby identifying potential outliers. We applied false discovery rate (FDR) correction using the R package *qvalue*, and we used 5% as the threshold for determining significant outlier traits. Results were visualized using Manhattan plots. The final set of outlier traits comprised the union of significant outliers detected across all three analyses. The outlier detection was conducted using the R package *pcadapt* ^*50*^.

## Data and Codes Availability

Data and codes sharing are not applicable until the manuscript has been accepted and published formally.

## Acknowledgements

This research was supported by the European Research Council (ERC) under the European Union’s Horizon 2020 research and innovation program (Grant Agreement No. 883700 TRAM), the National Natural Science Foundation of China (32470656 and T2122007), the National Key R&D Program of China (2020YFE0201600), the National Social Science Foundation (23&ZD317 and 20&ZD301), Shanghai Municipal Science and Technology Major Project (2017SHZDZX01). We also thank the Shanghai Institute for Mathematics and Interdisciplinary Sciences (SIMIS) for their financial support (grant number SIMIS-ID-2024-ZL). We thank the members of Linguistic Silk Road Research Consortium: Saiyinjiya Caidengduoerji, Siqinchaoketu, Yarjis Xueqing Zhong, Hao Li, Kai Sun, Jin Sun, Feiyiming Yan, Mingyuan Shao, Weibin Yin, Bingli Liu, Xingzhou Wei, Shuangcheng Wang, Mingyuan Shao, Wei Ma, and Zhencao Zhong for valuable assistance in data collection, collation, and annotation.

## Detailed member information in the Linguistic Silk Road Research Consortium

Department of English and Linguistics, Johannes Gutenberg University Mainz, Mainz, Germany

Dan Xu and Hao Li

Institute of Modern Languages and Linguistics, Fudan University, Shanghai, China

Menghan Zhang and Hongye Jin

Le Centre de recherches linguistiques sur l'Asie orientale (CRLAO), Paris, France

Saiyinjiya Caidengduoerji

Institute of Ethnology and Anthropology, Chinese Academy of Social Sciences, Beijing, China

Siqinchaoketu and Weibin Yin

School of Culture, History & Language, Australian National University, Canberra, Australia

Yarjis Xueqing Zhong

School of Liberal Arts, Nanjing University, Nanjing, China

Kai Sun

School of Sociology and Anthropology, Xiamen University, Xiamen, China

Jin Sun

Department of Chinese Language and Literature, Sun Yat-sen University, Guangzhou, China

Mingyuan Shao and Feiyiming Yan

Chinese Language and Culture College, Huaqiao University, Xiamen, China

Bingli Liu

School of Literature, Zhejiang University, Hangzhou, China

Xingzhou Wei

School of Humanities, Shanghai Normal University, Shanghai, China

Shuangcheng Wang

School of Literature and Communication, Qinghai Minzu University, Qinghai, China

Wei Ma

School of Chinese Ethnic Minority Languages and Literatures, Minzu University of China

Zhencao Zhong

